# Serratia marcescens Outer Membrane Vesicles rapidly paralyze Drosophila melanogaster through triggering apoptosis in the nervous system

**DOI:** 10.1101/2025.08.18.670980

**Authors:** Bechara Sina Rahme, Roberto E. Bruna, Marion Draheim, Chuping Cai, Maria Victoria Molino, Yaotang Wu, Miriam Wennida Yamba, Gisela Di Venanzio, Matthieu Lestradet, Eleonora García Véscovi, Dominique Ferrandon

**Affiliations:** Université de Strasbourg, CNRS, M3I UPR 9022 du CNRS, 67000 Strasbourg, France; Instituto de Biología Molecular y Cellular de Rosario, Consejo Nacional de Investigaciones Cientificas y Tecnológicas, and Facultad de Ciencias Bioquímicas y Farmacéuticas, Universidad Nacional de Rosario, Rosario S2000EZP, Argentina; Sino-French Hoffmann Institute, Guangzhou Medical University, Guangzhou 511436, China; Department of Molecular Microbiology, Washington University School of Medicine, St Louis, MO, 63110, USA

## Abstract

The pathogenicity of Gram-negative bacteria is mediated by multiple virulence factors that likely include secreted Outer Membrane Vesicles (OMVs) that can act as a cargo for delivery of enzymes or toxins to target tissues. Here, we have studied the effects on the host of OMVs prepared from one of the most potent pathogens of *Drosophila melanogaster*, *Serratia marcescens*. OMV injection leads to the apparent demise of flies within few hours. We identify a number of host defenses that somewhat protect it from the action of OMVs, namely the systemic humoral immunity pathway Immune deficiency, Prophenol Oxidases 1&2, and the redox active enzymes Dual oxidase, NADPH-oxidase, and Nitric Oxygen Synthase. In contrast, unidentified hemocyte function(s) and the circulating protease Hayan promote the pathogenicity of OMVs. Mechanistically, we find that OMVs promote the activation of the JNK pathway and the transient expression of the pro-apoptotic genes *head-involution defective* and *reaper* in at least neurons. Our data suggest that mitochondrially-derived reactive oxygen species promote neuronal cell death that leads to the paralysis of OMV-injected flies. We identify the metalloprotease PrtA as a major virulence factor of OMVs and show that the injection of purified PrtA mimics most of the effects of OMVs. Finally, our data further indicate that PrtA contributes to the pathogenicity of injected *Serratia marcescens*.

This study underscores the potential for OMVs to act as virulence factors that efficiently target the nervous system *in vivo* despite the blood brain barrier.

## Introduction

*Serratia marcescens*, a Gram-negative bacterium, belonging to Enterobacteriaceae, is found in diverse environments such as soil, water and air. *S. marcescens* has the capacity to infect plants, insects, and humans (Grimont and Grimont, 1978). As an opportunistic pathogen in humans, *S. marcescens* mainly causes nosocomial infections and is able to infect several human tissues such as the urinary (Marre et al., 1989), respiratory, endocardium or eye epithelia and may colonize the respiratory and digestive tracts (Albers et al., 2001). It is a health hazard in neonatal and adult intensive care units, in as much as it is often able to resist the action of multiple antibiotics and can be at the origin of septicemia (Hejazi and Falkiner, 1997). Generally, the pathogenicity of *S. marcescens* toward competing bacteria or to host cells, is mainly mediated through quorum sensing, the secretion of several virulence factors that include type-VI secretion systems (T6SS), a lipase, a phospholipase, a hemolysin, a DNase, the metalloprotease PrtA / Serralysin and related proteases, chitinases, and through the formation of outer membrane vesicles (English et al., 2012; Hejazi and Falkiner, 1997; Hertle, 2005; Lazzaro et al., 2017; McMahon et al., 2012; Murdoch et al., 2011; Petersen and Tisa, 2013; Shanks et al., 2015); furthermore, the virulence of clinical strains is highly correlated to the genetic inactivation of a two-component system required for T6SS activity that mediates a lifestyle switch that allows adaptation to a clinically-associated selection pressure (Williams et al., 2025).

Extracellular vesicles are produced and secreted by most living cells. For Gram-negative bacteria, these vesicles originate by a mechanism involving the pinching off of the outer membrane hence their name, outer membrane vesicles (OMVs). Besides incorporating outer membrane associated proteins, OMVs can encapsulate enzymes and virulence factors secreted to the periplasmic space, nucleic acids, and peptidoglycan fragments. Other secreted proteins can also rapidly associate with OMVs as documented for the full-length PrtV metalloprotease from *Vibrio cholerae* or the hemolysin from enterohemorrhagic *Escherichia coli* (Aldick et al., 2009; Rompikuntal et al., 2015). Their secretion helps bacteria to communicate with each other and mediate some of their interactions with the host (Goman et al., 2025; Kuehn and Kesty, 2005; Pin et al., 2023; Toyofuku et al., 2023). It has been demonstrated that OMVs participate in virulence by increasing bacteria communication and biofilm formation. They can be internalized by hosts in a variety of mechanisms such as endocytosis or fusion to lipid rafts. After internalization, OMVs or their cargo may escape to the cytosol by poorly described mechanisms that may in some cases be mediated by a pore-forming toxin (Bielaszewska et al., 2013). Several toxins delivered by OMVs have been documented to associate with mitochondria and to induce apoptosis (Bielaszewska et al., 2013; Deo et al., 2020; Deo et al., 2018). In addition, OMVs can protect virulence factors from host proteases, concentrate them prior to targeting them to host cells, and also contribute to their long-distance delivery (Bomberger et al., 2009; Toyofuku et al., 2023). A study of McMahon et al. (2012), reported that *S. marcescens* RM66262 (isolated from a human urinary tract infection (Bruna et al., 2015)) produced OMVs in a thermoregulated manner. Indeed, a decrease of the temperature (from 37 to 30 degrees) increased OMVs production without affecting their virulence since *G. mellonella* larvae succumbed at the same rate to OMVs from both temperatures (McMahon et al., 2012). A mass spectrometry analysis showed that virulence factors, such as the metalloprotease Serralysin (PrtA), are present in OMVs and not in the outer membrane compartment (McMahon et al., 2012).

*D. melanogaster* is a genetic invertebrate model organism, the host defense of which has been intensely investigated (Buchon et al., 2014; Lemaitre and Hoffmann, 2007; Liegeois and Ferrandon, 2022). The systemic humoral response against infections relies on two NF-κB signaling pathways. They mediate protection predominantly against fungal and Gram-positive infections (Toll pathway) or Gram-negative bacteria (Immune deficiency (IMD) pathway), essentially through the upregulation of antimicrobial peptide genes and other effectors. Melanization is an important host defense against microbial infections that is specific to invertebrates and it is mediated by cleaved pro-phenol oxidases, namely PPO1 and PPO2 in adults. Like Toll pathway extracellular activation, melanization depends on humoral proteolytic cascades, an important factor of which is the Hayan protease (Dudzic et al., 2019; Nam et al., 2012; Shan et al., 2023). Interestingly, Hayan is postulated to trigger a Hayan/PPO/ROS response, a cytoprotective program that defends neurons from traumatic injury (Nam et al., 2012). Finally, hemocytes, mostly plasmatocytes in adults, provide an additional arm of innate immunity, essentially but not only through phagocytosis (Melcarne et al., 2019; Vlisidou and Wood, 2015).

Host-pathogen interactions of *S. marcescens* with *Drosophila melanogaster* have been well-studied in both septic injury and oral infection models (Flyg et al., 1980; Kurz et al., 2003; Nehme et al., 2007; Lee et al., 2016; Sina Rahme et al., 2022; Socha et al., 2023). Among different *S. marcescens* strains, the Db10 strain has been isolated from moribund *Drosophila* flies and the Db10-derived Db11 strain is commonly used in *Drosophila* infection studies (Flyg et al., 1980). It was found that a few bacteria are sufficient to kill flies by bacteremia within 24h, making the Db10 strain one of the most potent *Drosophila* pathogens known to date (Nehme et al., 2007). *S. marcescens* RM66262 clinical strain displays similar virulence properties in *Drosophila* infection models (Sina Rahme et al., 2022). Although only few ingested bacteria manage to escape from the digestive tract within hours, they fail to rapidly kill the flies as they are controlled by plasmatocytes through phagocytosis (Cronin et al., 2009; Kocks et al., 2005; Nehme et al., 2007). Contrary to injected bacteria, the ingested bacteria fail to induce a systemic immune response but induce a local one in the midgut epithelium (Nehme et al., 2007).

In this work, we leverage our extensive knowledge of *Drosophila* innate immunity to investigate the mechanism underlying the virulence of OMVs, previously characterized in the insect *G. mellonella* (McMahon et al., 2012). Our findings reveal a complex interplay between host defenses and OMVs pathogenicity. Specifically, certain host factors, such as plasmatocytes and the Hayan serine-protease, enhance OMVs-induced pathogenesis, whereas others, including the IMD pathway, Dual Oxidase (Duox), and prophenoloxidases (PPOs), provide moderate protection. Furthermore, our data indicate that OMVs trigger a mitochondrial reactive oxygen species (ROS) response in neurons, leading to apoptosis within the nervous system. This neuronal damage accounts for the rapid demise of OMVs-injected flies, which is initially manifested as paralysis.

## Results

### The OMVs from *Serratia marcescens* are virulent to Drosophila

We injected wild-type flies with different concentrations of OMVs ranging from 0.01 to 0.1 ng/nL (Fig. 1A) and observed the demise of injected flies within hours, in a dose-dependent manner. The injection of 0.1 ng/nL OMVs led most of the flies to apparently succumb within 2 h, and lower concentrations led to a slower, less penetrant, lethality, whereas a 0.01 ng/nL concentration OMVs exhibited no virulence. We used a 0.1 ng/nL concentration for most experiments. Despite this, for each preparation OMVs concentration had to be finely adjusted due to batch-to-batch variations (that mostly depended on the length of storage of OMVs at -80°C).

**Figure 1:**
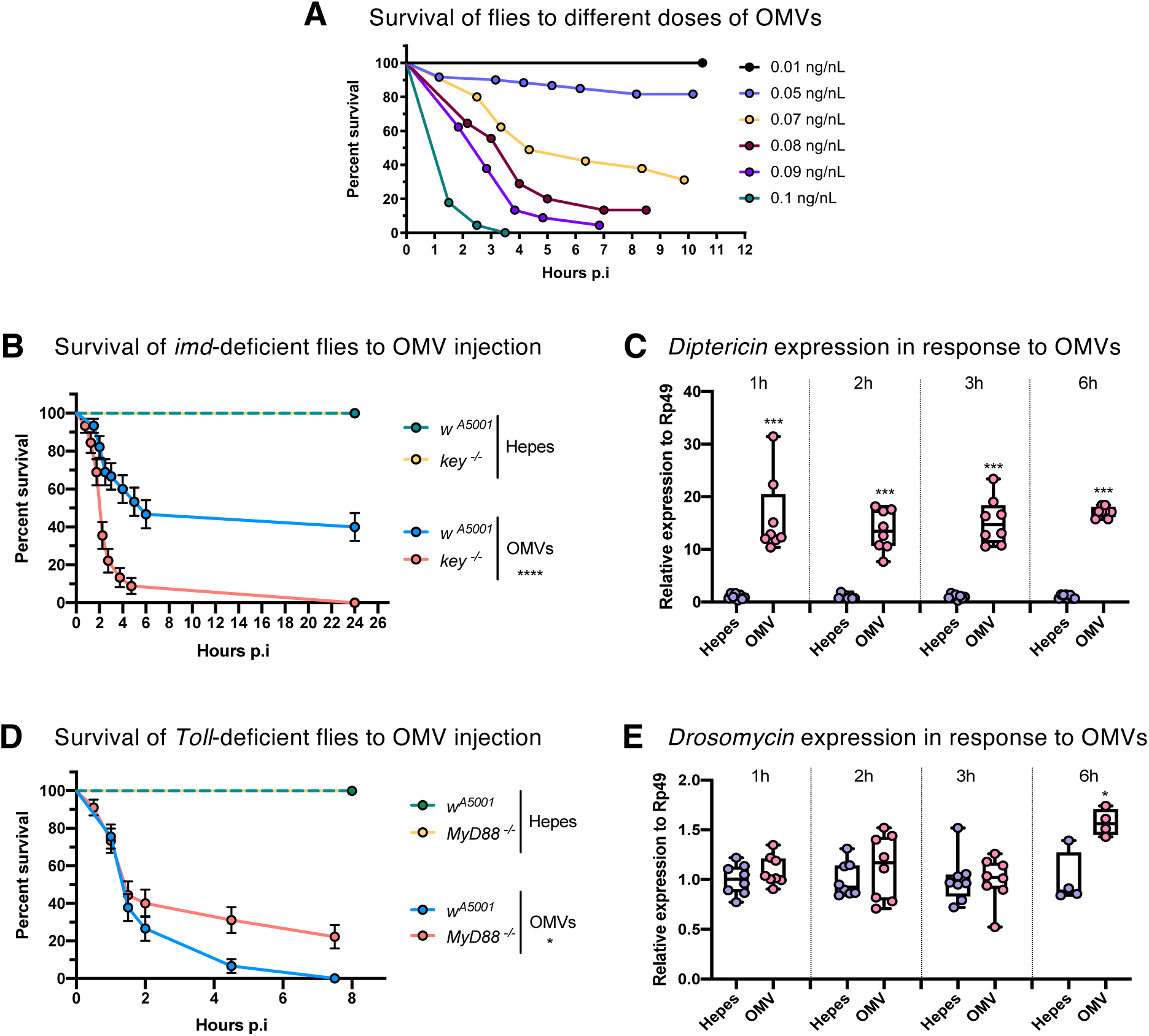
The IMD, but not the Toll pathway, is implicated in the host defense against *S. marcescens* OMVs. **(A)** Survival test of *w* [A5001] flies after the injection of different doses of purified *S. marcescens* OMVs. **(B, D)** Survival test of *key^-/-^* **(B)** and *MyD88^-/-^* **(D)** mutant flies to the injection of either 0.1 ng/μL of OMVs or HEPES buffer in comparison to wild-type flies (*w* [A5001]). Each graph represents one out of three independent experiments that yielded similar results. Error bars represent the standard error of the mean. Statistical tests were performed using the Logrank test. **(C, E)** RT-qPCR analysis of the *Diptericin* **(C)** and *Drosomycin* **(E)** expression in *w* [A5001] flies in response to OMV or HEPES buffer injection. Each dot represents the relative expression of the gene to *Rp49* (*RpL32*) of five adult females. Each graph represents the pooled data from three independent experiments. The middle bar of the box plots represents the median, and the upper and lower limits of boxes indicate the first and third quartiles, respectively; the whiskers define the minima and maxima. Statistical tests were performed using the Mann-Whitney test.

Upon a careful scrutiny, we noticed that flies that had fallen to the bottom of vials were not dead but paralyzed with some subtle and sporadic movements of the legs (Movie 1). Indeed, we observed pulsations of the dorsal vessel (Movie 2). Further, we found that the injected flies never recovered from this OMVs-induced paralysis that caused their ultimate death.

In conclusion, injected OMVs from *S. marcescens* exhibit high virulence in flies, resulting in apparent irreversible neurological damage.

### The IMD but not the Toll pathway is required for the host defense against OMVs

One of the cardinal features of *Serratia marcescens* infection is that injected bacteria appear to be insensitive to *Drosophila* host defenses, a property that requires an intact O-antigen (Kurz et al., 2003; Nehme et al., 2007). Thus, we tested the impact of the *Drosophila* systemic humoral immune response in the host defense against injected OMVs. Firstly, we monitored the survival rate of *D. melanogaster kenny* or *imd* null mutant strains that affect the IMD pathway. Compared to the wild-type strain, these mutants exhibited an increased sensitivity to the injection of OMVs (Fig. 1B, Fig. S1A). In keeping with these results, the expression of the IMD-dependent *Diptericin* gene was induced as early as 1 h post-injection (Fig. 1C). The situation for the Toll pathway appeared more complex. Specifically, a detectable induction of *Drosomycin*—used here as a readout for Toll pathway activation—was observed only at the 6 h post-injection time point, by which time the flies had already become paralyzed (Fig. 1E). Next, we challenged a set of Toll pathway mutants with OMVs injection. Most mutants, *MyD88*, *Toll, spätzle, GNBP3^hades^*and *Persephone-GNBP3^hades^* displayed a relatively delayed, and mildly enhanced resistance to injected OMVs, with *grass* showing a similar trend at a late phase of the experiment (Fig. 1D and Figs. S1B-F). *tube* and *pelle* showed survival rates comparable to the controls, whereas, unexpectedly, *Spätzle-processing enzyme* (*SPE*) mutants appeared more susceptible (Fig. S1G-I). *GNBP3* mutants exhibit an unexpected phenotype since *GNBP3* has been shown to encode a sensor for β-(1-3)-glucans, which are produced by fungi rather than bacteria (Mishima et al., 2009). However, there are reports of cyclic ß-(1-3)-glucans found in the bacterial periplasm that allow adaptation to osmotic stress (McIntosh et al., 2005). This raises the possibility that OMVs may carry such compounds that would be sensed by the GNBP3 sensor, for instance if OMVs are lysed through IMD pathway effectors. The opposite phenotypes displayed by *SPE* as compared to most Toll pathway mutants might suggest a noncanonical activation of the Toll pathway that deserves further in-depth studies. As regards Toll pathway activation, it might also be mediated by the PrtA protease (Bruna et al., 2018; El Chamy et al., 2008; Gottar et al., 2006; Issa et al., 2018). Nonetheless, the delayed activation of the Toll pathway observed after fly paralysis, combined with inconsistent phenotypic responses, suggests that the Toll pathway plays a limited role in the defense against injected OMVs, in contrast to the more prominent involvement of the IMD pathway.

### Plasmatocyte functions are required for the pathogenesis of OMVs

We next investigated whether the other arms of the innate immune response are involved in the host defense against OMVs. We first assessed the cellular immune response using three independent approaches.

We generated “hemoless” flies by inducing apoptosis in *hemolectin* expressing cells, which removes almost all phagocytic hemocytes (Charroux and Royet, 2009; Defaye et al., 2009). Unexpectedly, these flies were more resistant than control flies to OMVs challenge (Fig. 2A). Next, we saturated the phagocytic apparatus by prior injection of nondegradable “latex” beads (Lxb) and observed a similar, albeit somewhat weaker, resistance phenotype (Fig. 2B). Finally, we tested two null mutants for *eater*, which encodes a prospective phagocytosis receptor that also plays a role in the adhesion of hemocytes to tissues (Bretscher et al., 2015; Kocks et al., 2005). Again, a lesser susceptibility to injected OMVs was observed (Fig. S2A-B). Taken together, these observations suggest that the cellular immune response promotes the pathogenicity of injected OMVs.

**Figure 2:**
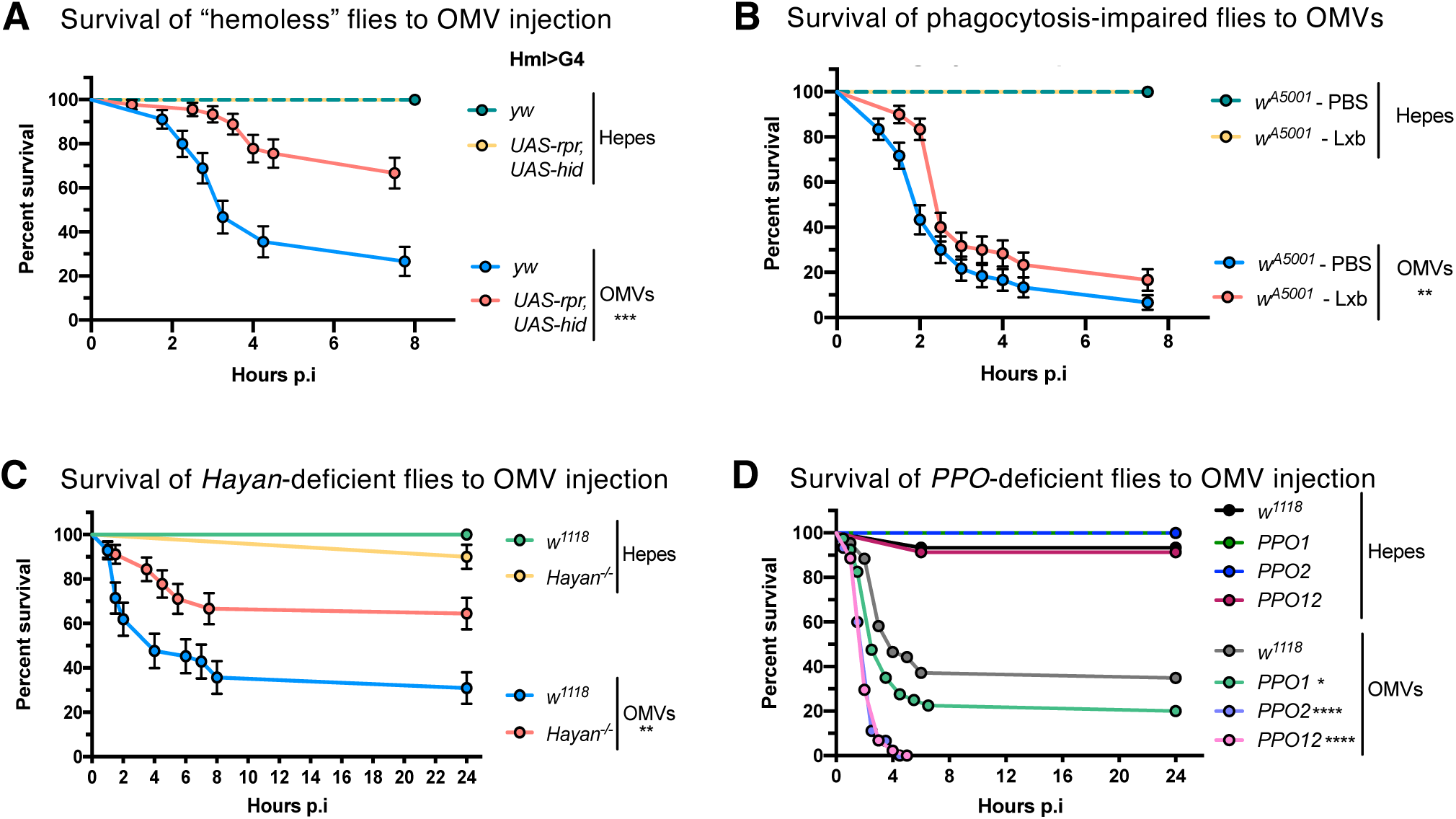
Phagocytes and *Hayan* promote the pathogenicity of *S. marcescens* OMVs, while PPOs play a protective role against OMVs. **(A)** Survival test of “hemoless” flies (*hmlG4>UAS-rpr, UAS-hid*) to the injection of either 0.1 ng/μL of OMVs or HEPES buffer in comparison to control flies (*hmlG4>yw*). **(B)** Survival test of phagocytosis-impaired flies injected with latex beads (Lxb) 24h prior to the injection of either 0.1 ng/μL of OMVs or HEPES buffer in comparison to flies that had been previously injected with PBS as a control. **(C-D)** Survival test of *Hayan^1^* **(C)** and *PPO1, PPO2* and *PPO1-PPO2* double mutants (*PPO12*) **(D)** flies to the injection of either 0.1 ng/μL of OMVs or HEPES buffer in comparison to wild-type flies (*w^1118^*). (A-D) Each graph represents one out of three independent experiments that yielded similar results. Error bars represent the standard error of the mean. Statistical tests were performed using Logrank.

### Hayan contributes to the pathogenesis of OMVs while PPOs are required in the host defense against OMVs

Next, we tested melanization, which is catalytically mediated by activated phenol oxidases that lead to the formation of melanin and also to a microbicidal activity that is possibly mediated by ROS (Dudzic et al., 2019; Moule et al., 2010; Nappi and Vass, 1993, 1996). A key player of the melanization response is the Hayan serine protease, which gets activated by the proteolytic cleavage of its N-terminal CLIP domain and, in *Drosophila,* is required for the cleavage of PPO to PO (Nam et al., 2012). We therefore monitored the survival rate of *Hayan* and *PPOs* deficient mutants upon OMVs injection. The loss of *Hayan* resulted in protection against OMVs (Fig. 2C, Fig. S2C). Surprisingly, *PPO1* and/or *PPO2* mutants were more susceptible to injected OMVs (Fig. 2D, Fig. S2D).

To our knowledge, it is the first time that opposite phenotypes are reported for *Hayan* and *PPO1/PPO2* mutants. In the Nam *et al*. model (Nam et al., 2012), it had been proposed that ROS are generated by POs activated first via Hayan. This is clearly not the case here as POs participate in the defense against injected OMVs whereas Hayan promotes their pathogenicity.

### ROS mediate the pathogenesis of OMVs

To examine the controversial results from melanization-related mutants, we directly investigated the role of reactive oxygen species (ROS) in the host response to injected OMVs. We determined that there is a significant increase in H_2_O_2_ levels 1 h after OMVs injection compared to buffer injection (Fig. 3A). To further evaluate the role of ROS, we co-injected OMVs with the vitamin C antioxidant. We added an additional control by pre-incubating OMVs with vitamin C followed by its removal using Amicon 10K filters. The co-exposure to vitamin C totally protected the flies from the action of OMVs whereas OMV pre-treatment with the antioxidant did not alter their virulence, suggesting that host-derived ROS play a significant role in mediating the pathogenicity of OMVs (Fig. 3B).

**Figure 3:**
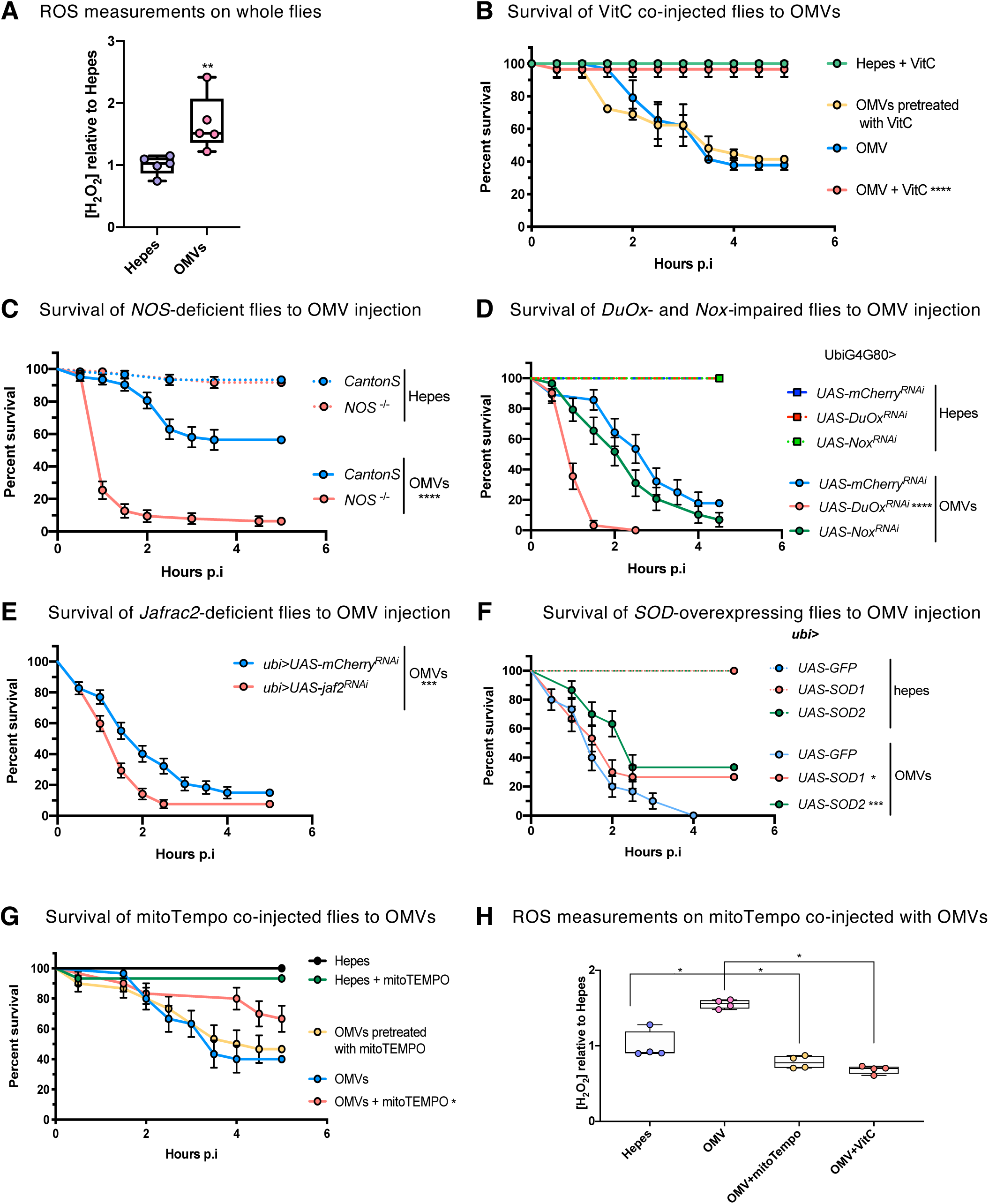
Mitochondrial ROS play a detrimental role in the stress response against *S. marcescens* OMVs. **(A)** Relative H_2_O_2_ measurements on single whole flies injected with 0.1 ng/μL of OMVs in comparison to HEPES injection. **(B)** Survival test of flies co-injected with 20 mM of Vitamin C and either 0.1 ng/μL of OMVs or HEPES in comparison to OMV only-injected flies. The yellow curve corresponds to OMVs that have been exposed to vitamin C for a limited period before its removal by filtration, prior to subsequent injection. **(C)** Survival test of *NOS*-deficient flies to OMV injection in comparison to *cantonS* control flies. (**D)** Survival test of *DuOx-* and *Nox*-impaired flies (*ubiG4G80>UAS-DuOx^RNAi^* and *ubiG4G80>UAS-NOX^RNAi^*) to the injection of OMVs in comparison to control flies (*ubiG4G80>mcherry^RNAi^*). **(E)** Survival test of *Jafrac*-deficient flies (*ubiG4>UAS-jaf2^RNAi^*) to the injection of 0.1 ng/μL of OMVs in comparison to control flies (*ubiG4>mcherry^RNAi^*). **(F)** Survival test of *SOD*-overexpressing flies (*ubiG4>UAS-SOD1* and *ubiG4>UAS-SOD2*) to the injection of 0.1 ng/μL of OMVs in comparison to control flies (*ubiG4>UAS-GFP*). **(G)** Survival test of flies co-injected with 20 μM of mitoTEMPO and either 0.1 ng/μL of OMVs or HEPES buffer in comparison to HEPES- or OMV-injected flies. The yellow curve corresponds to OMVs that have been exposed to mitoTEMPO for a limited period before its removal by filtration, prior to subsequent injection. **(H)** Relative H_2_O_2_ measurements on single whole fly co-injected with 20 μM of mitoTEMPO or vitaminC and either 0.1 ng/μL of OMVs or HEPES buffer in comparison to HEPES and OMV injection. (A-H) Each graph represents one out of three independent experiments that yielded similar results. (B-G) Statistical tests were performed using Logrank, and error bars represent the standard error of the mean. (A, H) The middle bar of the box plots represents the median, and the upper and lower limits of boxes indicate the first and third quartiles, respectively; the whiskers define the minima and maxima. Statistical tests were performed using Mann-Whitney.

Next, we aimed to assess the source of host-generated ROS using a genetic approach. A nitric oxide synthase null mutant (*NOS*) or the ubiquitous silencing of the *Nox* and *Duox* genes revealed that *NOS* and *Duox* are involved in the defense against injected OMVs and not in promoting their pathogenicity (Fig. 3C-D). *Jafrac2* acts intracellularly as it encodes a peroxiredoxin located in the endoplasmic reticulum. *Jafrac2* silencing in all tissues also revealed its involvement in the host defense against OMVs (Fig. 3E).

Another potential source of intracellular ROS is constituted by mitochondria. ROS released from the electron respiratory chains are transformed into H_2_O_2_ by superoxide dismutases (SOD). The ubiquitous overexpression of a mitochondrial SOD gene, *SOD2*, yielded a phenotype of enhanced resistance to injected OMVs, suggesting that mitochondria may mediate the noxious effects of ROS upon OMVs injection. In contrast, the overexpression of a cytoplasmic SOD gene, *SOD1*, provided a milder degree of protection (Fig. 3F). To independently confirm these results, we co-injected OMVs alongside a strong, mitochondrially-targeted, antioxidant, Mito-TEMPO (Oyewole and Birch-Machin, 2015). Results similar to those obtained with Vitamin C were obtained, namely that the co-injection protected the host from the harmful effects of mitochondria-generated ROS while pre-incubation of mito-TEMPO with OMVs retained their virulence. We also observed that the use of mito-TEMPO inhibited the production of hydrogen peroxide, which suggests that the peak of production observed in wild-type flies is essentially produced by mitochondria (Fig. 3G-H).

These results indicate that mitochondria-generated ROS contribute significantly to the pathogenicity of injected OMVs. Conversely, other ROS, likely secreted, appear to play a protective role by defending against OMVs.

### JNK pathway activation in neurons is detrimental to the host upon OMVs challenge

The JNK pathway is known to be induced upon ROS stress and also to activate apoptosis. In light of our results, we investigated whether the JNK pathway might become activated upon OMVs injection. *Puckered* encodes a phosphatase that acts as a negative regulator of the JNK pathway and is often used as a JNK pathway activation read-out. *Puckered* mRNA levels were induced 1 h after OMVs challenge (Fig. 4A). We therefore ubiquitously silenced *kayak*, which encodes cFos, a component of the AP1 transcription factor that mediates much of JNK pathway activation, and observed an increased resistance to OMVs in “survival” experiments (Fig. S3A). Given the paralysis phenotype, we next asked whether JNK activation in neurons might be promoting the pathogenicity of OMVs. Upon silencing *kayak* in neural cells using an *elav-Gal4* driver, we observed a strong, reproducible protection against injected OMVs (Fig. 4B). In contrast, silencing *kayak* in *hml-*positive hemocytes yielded a slight susceptibility to injected OMVs (Fig. S3B). Of note, we observed a transient induction of the JNK pathway in head and thorax tissues, which contain most of the fly nervous system and also the cephalic fat body (Fig. S3C).

**Figure 4:**
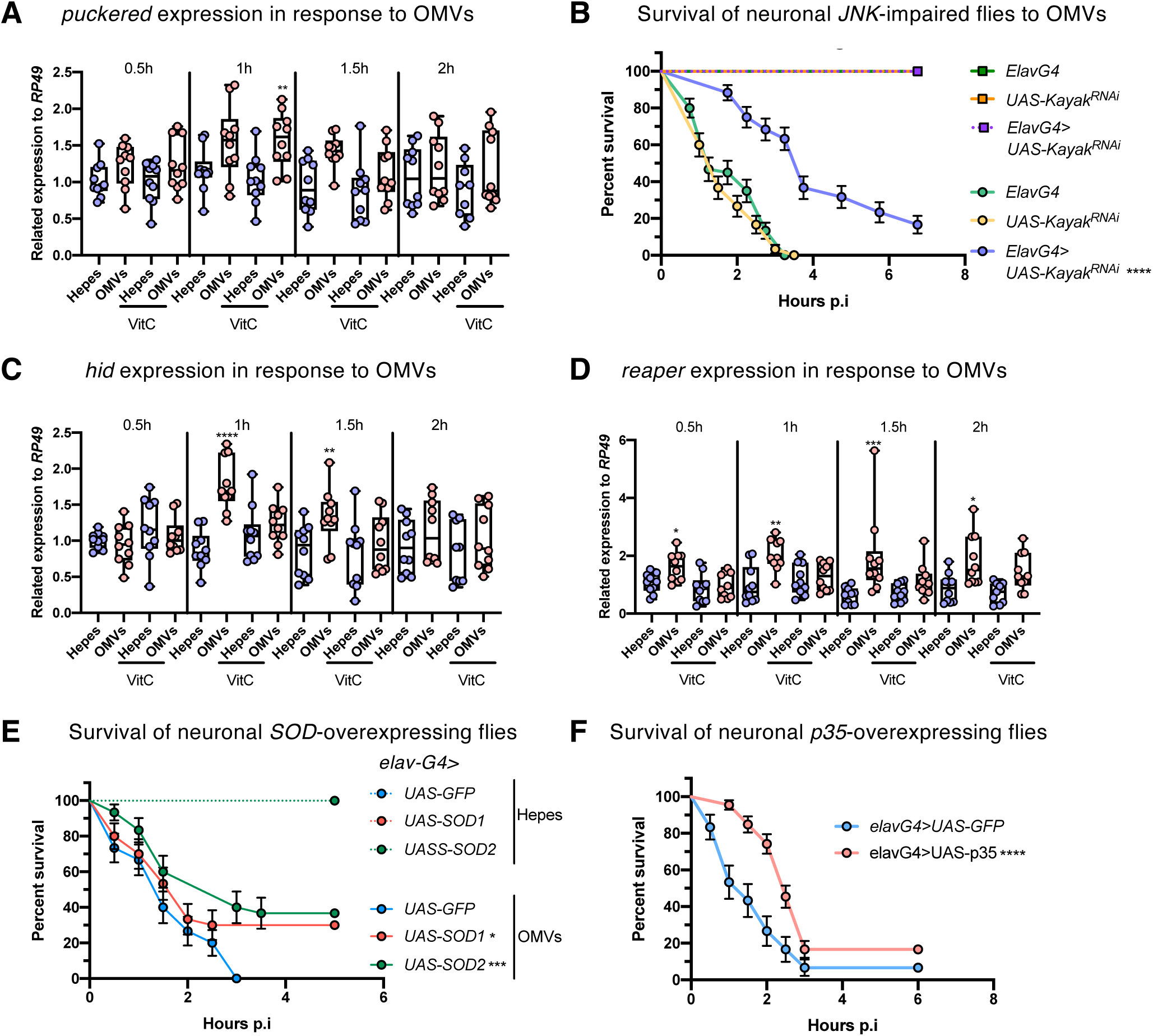
The neuronal JNK pathway promotes the pathogenicity of *S. marcescens* OMVs **(A, C-D)** RT-qPCR analysis of *puckered*. **(A)***, hid* **(C)** and *reaper* **(D)** expression in *w* [A5001] flies in response to the OMV or HEPES buffer injection or in response to the co-injection of 20 mM of Vitamin C with either OMV or HEPES at 0.5, 1, 1.5 and 2 h after injection. Each dot represents the relative expression of the gene to *Rp49* (*RpL32*) of 5 adult females. The expression of each gene for each condition is relative to its expression in the HEPES-injected flies. Each graph represents the pooled data from two independent experiments. The middle bar of the box plots represents the median, and the upper and lower limits of boxes indicate the first and third quartiles, respectively; the whiskers define the minima and maxima. Statistical tests were performed using Mann-Whitney. **(B)** Survival test of flies in which *JNK* expression has been silenced by RNAi specifically in neurons (*elavG4>UAS-kayak^RNAi^*) to the injection of either 0.1 ng/μL of OMVs or HEPES buffer in comparison to control flies (*elav-G4* and *UAS-kayak^RNAi^*). **(E)** Survival test of flies in which *SOD* genes are overexpressed in neurons (*elavG4>UAS-SOD1* and *elavG4>UAS-SOD2*) to the injection of either OMVs or HEPES buffer in comparison to control flies (*elavG4>UAS-GFP*). **(F)** Survival test of flies in which the baculovirus *p35* gene is overexpressed in neurons (*elavG4>UAS-p35*) to the injection of OMVs in comparison to control flies (*elavG4>UAS-GFP*). (B, E & F) Each graph represents one out of three independent experiments that yielded similar results. Statistical tests were performed using Logrank, and error bars represent the standard error of the mean.

### Apoptosis contributes to the pathogenesis of OMVs in Drosophila

JNK pathway activation is known to induce apoptosis through the up-regulation of the expression of the inhibitor of apoptosis (IAP) antagonists *reaper* (*rpr*) and *head involution defective* (*hid*) genes; thus, we monitored their induction using RT-qPCR. Interestingly, *hid* expression was induced 1 h and 1.5 h whereas *rpr* induction was detected between 0.5 to 2h after OMVs challenge (Fig. 4C-D). We next assessed whether ROS might be required for the induction of *rpr* and *hid* expression. Upon vitamin C co-injection of OMVs, the induction of these genes was abolished (Fig. 4C-D). Unexpectedly, under these conditions, *puckered* was still induced 1 h post inoculation (Fig. 4A).

In keeping with these results, we next overexpressed *SOD* genes in neurons and observed that *SOD2* ectopic expression in neurons did protect the flies from injected OMVs pathogenicity (Fig. 4E, Fig. S3D).

The apoptosis program involves executioner caspases that can be inhibited by the baculovirus p35 effector (Hay et al., 1994). We therefore ectopically expressed p35 in neurons using either *elav-Gal4* or *elav-GS (*Gene Switch) drivers, which can be activated upon administration of RU486 to flies. In both cases, the *p35*-overexpressing flies displayed an enhanced resistance phenotype (Fig. 4F, Fig. S3E). Taken together, these data suggest that OMVs induce apoptosis in some neurons through the generation of ROS, an effect that significantly contributes to their pathogenicity.

### PrtA is a major virulence factor of OMVs

A previous proteomic analysis of OMVs content revealed that the 56kD metalloprotease PrtA, also known as Serralysin, is packaged into OMVs (McMahon et al., 2012). We thus prepared OMVs from a *prtA* mutant strain and injected the preparation into flies. *PrtA*-deficient-OMVs (*prtA*-OMVs) were avirulent when injected at our usual concentration and flies were not paralyzed (Movie 3). Nevertheless, they displayed a dose-dependent pathogenicity when injected at 10 to 100x higher concentrations (Fig. 5A-B). Because the lack of PrtA might not be the only alteration of the OMVs composition when obtained from the *S. marcescens prtA* mutant strain, and to dissect the role of PrtA, we injected purified PrtA. We used a concentration estimated to be equivalent to the one present in OMVs at the 0.1 ng/nL concentration that also yielded an equivalent demise of injected flies (Fig. 5C). The concomitant injection of purified PrtA was able to rescue the decreased virulence of *prtA*-OMVs (Fig. 5D).

**Figure 5:**
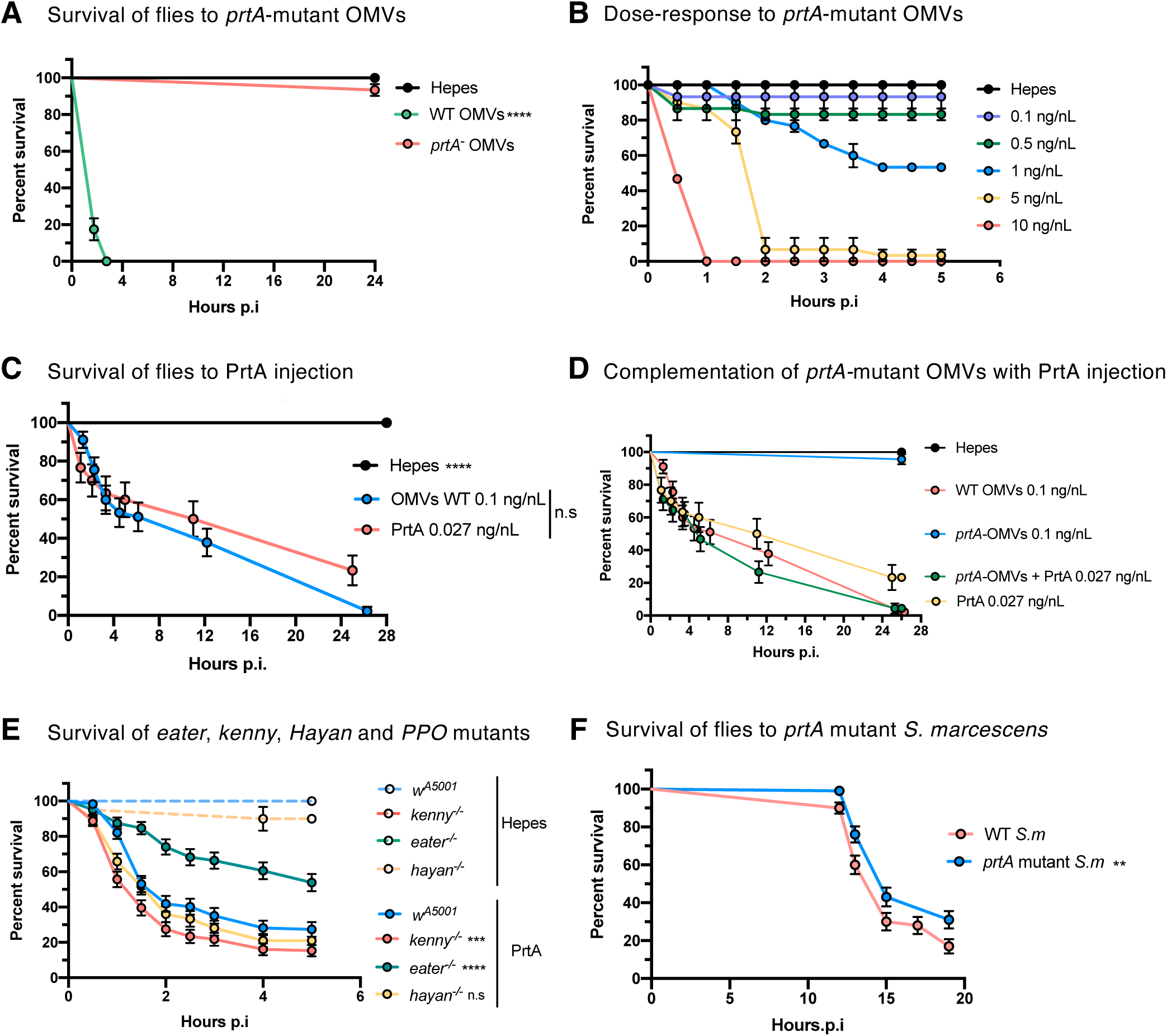
The virulence of *S. marcescens* OMVs is mediated by the PrtA metalloprotease. **(A**) Survival test of *w* [A5001] flies after the injection of 0.1 ng/μL of OMVs purified from either WT (WT OMVs) or *prtA* mutant (*prtA^-^* OMVs*) S. marcescens.* **(B)** Survival test of *w* [A5001] flies after the injection of different doses of OMVs purified from *prtA* mutant *S. marcescens.* **(C)** Survival test of *w* [A5001] flies after the injection of 0.1 ng/μL of OMVs or 0.025 ng/μL of purified PrtA protease from WT *S. marcescens*. **(D)** Survival test of *w* [A5001] flies after the co-injection of 0.1 ng/μL of *prtA* mutant OMVs and 0.025 ng/μL of purified PrtA (*prtA^-^* OMVs complementation) in comparison to flies injected with 0.1 ng/μL of WT OMVs. **(E)** Survival test of *kenny, eater* and *Hayan* mutants to the injection of 0.27 ng/μL of purified PrtA in comparison to HEPES injection and *w* [A5001] control flies. **(F)** Survival test of *w* [A5001] flies after the injection of *prtA* mutant *S. marcescens* bacteria in comparison to the injection of WT RM66262 bacteria. (A-F) Each graph represents one out of three independent experiments that yielded similar results. Statistical tests were performed using Logrank, and error bars represent the standard error.

To assess whether PrtA might act in a manner similar to that of injected wild-type OMVs, we injected purified PrtA into wild-type and immunodeficient flies. Whereas *eater* and *kenny* mutants exhibited a phenotype similar to that observed upon the injection of OMVs, *i.e.*, respectively an enhanced resistance or sensitivity (albeit mild as compared to that observed with OMVs), it was striking that *Hayan* mutant flies did not display any altered behavior in the “survival” experiment (Fig. 5E). This observation suggests that Hayan may be required to activate the proteolytic activity of PrtA embedded in OMVs.

Finally, we investigated the role of OMVs in the high virulence of *S. marcescens* using a septic injury model. Because PrtA is a key mediator of OMVs virulence, we hypothesized that if OMVs significantly contribute to the bacterium’s virulence in vivo, then a *prtA* mutant strain would exhibit reduced virulence. Consistently, we observed a mild but statistically significant reduction in the pathogenicity of the injected *prtA* mutant bacteria compared to the wild-type strain (Fig. 5F).

Given that pathogens typically rely on multiple virulence factors, this outcome was not unexpected. It shows that PrtA contributes to the virulence of *S. marcescens* in the septic injury model, potentially by reaching pathogenic levels when bacterial titers, and therefore the release of OMVs, are sufficiently high.

## Discussion

*S. marcescens* is one of the most potent pathogens of *Drosophila* in a systemic infection model since the injection of very few bacteria (<10) causes the death of flies in less than a day (Nehme et al., 2007). In this work, we have established that purified *S. marcescens* OMVs lead to the demise of injected flies within hours, provided they are injected at a sufficiently high concentration that might be reached toward the end of *in vivo S. marcescens* infection. An important observation was that flies were actually not killed but were initially paralyzed, suggesting an action of OMVs on the neuro-muscular system. We have shown that the different arms of the innate immune system play contrasted roles in the host defense against injected OMVs. Whereas the IMD pathway, phenol oxidases, and some extracellular ROS-generating enzymes such as NOS or Duox decrease the pathogenesis of the injected OMVs, others, notably the cellular immune response and the Hayan protease, promote it. We further establish that the injection of OMVs leads to the production of harmful mitochondrial ROS that trigger apoptosis in neurons possibly through the activation of the JNK pathway.

The IMD pathway fosters a defense against injected OMVs, in keeping with its general role in fighting off Gram-negative bacterial infections (Lemaitre and Hoffmann, 2007). One may envision that membrane active AMPs such as Cecropins might lyse OMVs, thereby leading to the release of OMV content in the hemolymph. However, these peptides are unlikely to be active against injected purified PrtA, which kills IMD pathway mutant flies faster than wild-type flies. We have recently identified two peptidic effectors of the systemic immune response, Shenshu and Yulü, the expressions of which are dependent on the IMD pathway, that are able to counteract the noxious activity of PrtA, without directly inhibiting this metalloprotease catalytic activity in *in vitro* assays (Cai *et al*., *in revision*). Other IMD-regulated effectors such as *Drosophila* complement protein family members of *Drosophila* do further contribute to the host defense against PrtA and OMVs through their α2-macroglobulin-like, Shenshu/Yulü-dependent, inhibition of PrtA catalytic activity (Cai *et al*., *in revision*). These peptides are however likely to mediate only part of the IMD-pathway mediated host defense against *S. marcescens*RM66262 since the *shenshu-yulü* double mutant has a less pronounced sensitivity phenotype as *kenny* mutants (Cai *et al*., *in revision*).

The apparently paradoxical role of the melanization activation cascade is puzzling at first sight. However, an earlier study has revealed an uncoupling between the Hayan-dependent synthesis and deposition of melanin at the wounding site and a killing activity mediated by the Sp7 protease in the host defense against low doses of injected *Staphylococcus aureus* bacteria (Dudzic et al., 2019). One may thus envision that Sp7 activates PPO1 and PPO2 generating an activity that mitigates the pathogenic effects of OMVs—possibly through ROS production, in keeping with those already potentially generated by NOS, Duox or other metabolites. In contrast, the exact function of Hayan in promoting the pathogenicity of OMVs but not of injected PrtA remains to be exactly delineated; our data are compatible with Hayan acting and activating PrtA present in OMVs. The potential relationship of Hayan to plasmatocytes that are also required for the detrimental effects of injected OMVs requires further clarifications. We did find that Eater also promotes the pathogenicity of injected PrtA and not only that of OMVs, unlike Hayan.

As our data document an involvement of the nervous system in mediating the noxious effects of OMVs, an interesting issue is whether OMVs can actually access the brain and have the ability to cross the blood-brain barrier (BBB). Our attempts to label *S. marcescens* OMVs by tagging one of its constituents, PrtA, have failed and we have not been able to directly assess their presence in this tissue using microscopy techniques. The OMVs from *Porphyromonas gingivalis* have been reported to weaken the BBB likely through the digestion of tight junction proteins by gingipains, cysteine proteases associated with *P. gingivalis* OMVs (Nonaka et al., 2022; Pritchard et al., 2022). Thus, it is an open possibility that OMVs might reach and act on neurons.

The model we propose to account for our data is that injected OMVs trigger the production of ROS by mitochondria in neurons, which in turn would cause apoptosis. Several studies have documented that OMVs can damage mitochondria through different virulence factors (Bielaszewska et al., 2013; Deo et al., 2020; Deo et al., 2018; Tiku et al., 2021). In the case of *Acinetobacter baumannii*, the OmpA porin induces mitochondrial fragmentation, which correlates also with mitochondrial damage as evidenced by reduced ATP levels and depolarization of mitochondria and the production of reactive oxygen species (ROS) (Tiku et al., 2021). The JNK pathway is known to be triggered by ROS and to promote the expression of *rpr* and *hid,* yet we observed that it was still induced in vitamin C-injected flies, for which *rpr* and *hid* induction was impaired. Thus, it may be that the JNK pathway and Rpr and Hid function in parallel in triggering apoptosis. Indeed, both pro-apoptotic proteins become associated with the mitochondria outer membrane where they would trigger apoptosis by inhibiting the apoptosis inhibitor DIAP-1 (Clavier et al., 2016). It will be important to determine whether apoptosis in the brain occurs in a random widespread manner or if it targets neuronal specific circuits that regulate locomotion and posture. Therefore, OMVs may cause fly mortality either through combined effects on multiple organs or, alternatively, by causing damage primarily confined to the brain, ultimately leading to death through starvation as flies are paralyzed.

Finally, PrtA/serralysin is used as a therapeutic enzyme under the denomination serratiopeptidase because of multiple beneficial effects in a number of medical conditions, having been approved as a nutritional supplement or an active pharmaceutical ingredient in a number of countries following various *in vitro, in cello*, or clinical studies (Hosseini et al., 2024). It had been originally isolated from enteric Serratia species found in the silkworm in the late 60’s (Yamasaki et al., 1967). Among its reported activities are anti-inflammatory, anti-edema, and anti-biofilm properties as well as its ability to dissolve dead tissues, clots, amyloid fibers, arterial plaque while not attacking live tissues. Remarkably, adverse mild to moderate side-effects have been rarely reported. This family of proteases is undergoing several improvements to increase its efficiency as well as its delivery: it is mostly absorbed through oral administration, which drastically decreases its bio-availability to amounts that end up being much lower to those used in this study. Thus, PrtA being before all a virulence factor, care should be taken to remain in a safety concentration/efficiency range to avoid potential effects on the nervous system.

## Material and Methods

### Bacterial culture

*S. marcescens* RM66262 (Bruna et al., 2015) and the otherwise isogenic *prtA* mutant strains were cultured on Lysogeny Broth (LB)-agar plates containing ampicillin or chloramphenicol antibiotic. Bacteria were grown overnight on the agar plates at 37°C. These plates were stored at 4°C for one week. One colony was inoculated in liquid LB and the culture was kept at 30°C with agitation for a maximum of 16 h for OMVs production.

### Purification of outer membrane vesicles (OMVs)

Saturated cultures of RM66262 or *prtA* mutant were diluted 1:500 in fresh LB and incubated at 30°C with agitation for a maximum of 16 h. About 600 mL of cultures were centrifuged at 8,000 g for 15 min. The pellets were discarded, and the supernatants were filtered using 0.45 μm filter (Thermo scientific, cat# 126-0045) to remove all remaining cells. The filtered supernatants were further precipitated with ammonium sulfate (129g/250mL of supernatant) overnight at 4°C. Then, the supernatants were centrifuged for 20 min at 10,000 g. The supernatants were discarded, and the pellets were re-suspended in 10 mL of 50 mM HEPES buffer (pH 7.5). The samples were dialyzed against HEPES (50 mM) overnight at 4°C, then concentrated using Amicon® Ultra 15 mL (Millipore, UFC901024) by centrifugation at 5,000 g. Further, 4 mg from the samples were loaded on the top of a 30% to 60% sucrose/HEPES density gradient and centrifuged at 100,000 g at 4°C for 16 h. Fractions of 1 mL were collected from each gradient and checked on Coomassie gel. Only the 6 first fractions were selected and dialyzed against HEPES 50 mM overnight at 4°C. Finally, the samples were centrifuged at 100,000 g for 3 h at 4°C, the supernatants were discarded and the pellets that contain OMVs were re-suspended in 400 μL of fresh 50 mM HEPES (pH 7.5). The OMVs were stored at -80°C.

### Fly strains and maintenance

Fly lines were raised on media at 25℃ with 65% humidity. For 25 L of fly food medium, 1.2 kg cornmeal (Priméal), 1.2 kg glucose (Tereos Syral), 1.5 kg yeast (Bio Springer), 90 g nipagin (VWR Chemicals) were diluted into 350 mL ethanol (Sigma-Aldrich), 120 g agar-agar (Sobigel) and water qsp were used.

Fly strains used in the experiments were: control flies *w* [A5001], *w^1118^*, *yw*, GD line, KK line or mCherry as indicated, *eater*^-/-^ mutant flies (Bretscher et al., 2015; Kocks et al., 2005), *MyD88*^-/-^, *Hayan^1^* (Nam et al., 2012), *Hayan^34^* (BL 67097), *PPO1*^-/-^ and *PPO1-PPO2* ^-/-^ (Dudzic et al., 2015), *imd^shadok^* (Gottar et al., 2002). The RNAi lines used were from the Vienna *Drosophila* Research Center (VDRC) or were Bloomington TRiP lines (BL):, UAS-*kayak^RNAi^* (VDRC KK #19512), UAS-*Jafrac2 ^RNAi^* (TRiP lines, TH03349.N), UAS-*DUOX^RNAi^* (BL# 38907). *NOS* null mutants were a kind gift from Dr. Patrick O’Farrell (Yakubovich et al., 2010). *ubiGal4, Gal80^ts^>UAS-transgene* crosses were performed at 18°C. The progeny was kept at 18°C until hatching and was transferred to 29°C for five days prior to experiments.

### Injections of OMVs and treatment

The injection of OMVs into the thorax of the flies was carried out with a Nanoject II auto-nanoliter injector (Drummond). Depending on the OMVs batch 69 nL of 0.07 to 0.1 ng/nL of OMVs were injected. The Vitamin C antioxidant treatment or mito TEMPO treatment (SML0737) were performed through a co-injection of OMVs (0.1 ng/nL) with Vitamin C (20 mM) or mito TEMPO (20µM). All OMV samples and antioxidants were diluted in 50 mM HEPES (pH 7.5). For pre-treatment, OMVs were incubated with or without Vitamin C (20mM) or mito TEMPO (20µM) for 30mn on ice. OMVs were centrifuged 15mn 10,000g using a 10kD Amicon filter to remove the chemical from the solution prior to OMV injection.

### Purification of PrtA

A *S. marcescens slpE* mutant strain was used as a production microorganism since the molecular weight and isoelectric point of the SlpE metalloprotease is similar to PrtA and could interfere with the purification of native PrtA. Cells were grown at 30 °C for 16 h in SLB medium (10 g/L tryptone, 5 g/L yeast extract), centrifuged and the recovered supernatant was filtered and concentrated with a Centricon. The supernatant was applied to an anion exchange column (Mono Q) and elution was carried out by a linear NaCl gradient. The active fractions were collected, and concentration and proteolytic activity were measured (using azocasein as substrate), to ensure that the protease is catalytically active after purification.

### Survival tests to OMV injections

Survival tests were performed using 15-20 flies per vial in biological duplicates or triplicates. The number of flies alive was counted every 30 min after the injection. Of note, only flies that completely stopped moving their legs were considered to be dead.

### Quantification of gene expression

Four to five whole flies were crushed into 100 μL of Trizol. Samples were filled with 900μl of Trizol and mixed with 500 µl of chloroform. Samples were centrifuged at 12,000 g for 15 min at 4°C. The liquid upper phase of the samples was collected into an Eppendorf tube containing 100 μL of isopropanol. The tubes were vortexed and incubated at room temperature for 10 min. The samples were then centrifuged at 12,000 g for 15 min at 4°C. The pellet was washed in 500 μL of 70% ethanol and dried. RNAs were then re-suspended in DEPC water. A volume of 10 μL was used to generate cDNA by reverse transcription, using the Transcript II all in one first strand synthesis supermix for qPCR (one step gDNA removal) synthesis kit (transgene biotech #AT341-02). The quantitative Polymerase Chain Reaction (qPCR) was performed with the same kit on cDNA diluted 20 times. The program-used was the following: 30 sec at 98°C; 34 cycles of 5 sec at 95°C, 30 sec at 98°C and finally 30 sec at 65°C. The data were analyzed using the CFX384 software (Bio-Rad). The Ct (Cycle threshold) values of the genes were normalized with the Ct values of Rpl32 (housekeeping gene that codes for ribosomal protein rp49). Relative gene expression levels were calculated using the ΔΔCt method. The following primer couples were used for RTqPCR:

*Drosomycin* Fwd: 5’ CGTGAGAACCTTTTCCAATATGATG 3’

*Drosomycin* Rev: 5’ TCCCAGGACCACCAGCAT 3’

*Dpt* Fwd: 5’ GCTGCGCAATCGCTTCTACT 3’

*Dpt* Rev: 5’ TGGTGGAGTGGGCTTCATG 3’

*puc* Fwd: 5’ GCTGAACGTTACCTGCCAAA 3’

puc Rev: 5’ TTGCATGTACTTGAGGCCCT 3’

### H_2_O_2_ measurement

Briefly, single flies were homogenized in the specific buffer provided by Sigma Aldrich kit MAK165. Then, 10µL of each sample were used to measure H_2_O_2_ concentrations following the protocol provided by the manufacturer.

### Western blot

For western blots, hemolymph samples were collected from 50 flies in a protease inhibitor cocktails solution. Protein concentration of the samples was determined by Bradford assay. 30 µg protein was separated on an 8% gel by SDS-PAGE and transferred to a PVDF membrane. After blocking in 5% bovine serum albumin in PBST for 1 h at room temperature, samples were incubated at 4 °C overnight with rabbit antibodies against *Drosophila* PPO1 at a 1:10,000 dilution (a kind gift from Prof. Erjun Ling). After washes, a goat anti-rabbit-horseradish peroxidase (HRP) secondary antibody at a 1:20,000 dilution was incubated for 1 h at room temperature. Enhanced chemiluminescence substrate (ECL, General Electric Healthcare) was used to reveal the blot according to the manufacturer’s instructions.

### Statistical tests

All graphs and statistical tests were analyzed using the GraphPad software Prism 6 or 8. The statistical test used for the survival tests was Logrank. The Mann-Whitney test was performed on all other experiments. The number of stars represents the P values P≥0.05 (ns), P<0.05 (*), P<0.01 (**), P<0.001 (***) and P<0.0001 (****).

## Acknowledgments

We are grateful to J. Nguyen for help in some experiments; to Drs Bruno Lemaitre, David Gubb, Won-Jae Lee, Patrick O’Farrell and to the VDRC and Bloomington Stock Centers (NIH P40OD018537) for fly stocks, to Prof. Erjun Ling for antibodies. G.D.V RB, and VM have been supported by ANPCyT (Agencia Nacional de Promoción Científica y Tecnológica, Argentina) and CONICET (Consejo Nacional de Investigaciones Cientifícas y Técnicas, Argentina), and B.S.R by an IdEx fellowship (Université de Strasbourg-France). The work in E.G.V laboratory has been funded by CONICET (Argentina) and the work in D.F laboratory has been funded by CNRS (Centre National De La Recherche Scientifique), Fondation pour la Recherche Médicale (Equipe FRM to DF [FRM DEQ20090515394]), and ANR (Agence Nationale de la Recherche, France, [ANR-11-EQPX-0022, DROSOGUT, ENTEROCYTE_PURGE_RECOVERY]). The D.F and E.G.V laboratories have developed a collaboration within the framework of the ECOS-Sud (France)-MINCT (Ministerio deCiencia y Tecnología e Innovación) exchange program (#A12B04). MD and CC have been supported by Guangzhou Medical University; CC has also been supported by the China Scholarship Council (#202108440349). Work in Guangzhou Medical University was also supported by grants from the 111 Project (#D18010; China), the Incubation Project for Innovative Teams of the Guangzhou Medical University, the Open Project from State Key Laboratory of Respiratory Diseases, China, and the China High-end Foreign Talent Program to DF.

